# Small molecules from Bacopa monneiri as potent inhibitors against Neurodegenerative disorders

**DOI:** 10.1101/2022.03.31.486590

**Authors:** Satyam Sangeet, Arshad Khan

## Abstract

Alzheimer’s is characterized by the formation of senile plaques and fibril tangles. Several methodologies have been employed to treat the disease. Albeit engineered medications which are accessible for the treatment of Alzheimer’s, due to their numerous side-effects, it becomes imperative to formulate and synthesize novel drug candidates. Plants could be utilized as an alternative for these manufactured medications because of their low incidental effects in contrast with the engineered drugs. *Bacopa monneiri* (BM) is a therapeutic plant which is accounted for to be utilized to treat NDs. Therefore, in current study an *in-silico* approach was carried out to evaluate the pharmacological effect of BM. Molecular Docking was carried out to screen the active phytochemicals of BM which can act as potential drug candidates against amyloid-β plaques. A total of 8 biologically active phytochemicals from BM were docked against p75NTR receptor. Based on molecular docking study it was observed that the phytocompounds Bacopasaponin D and Bacopasaponin G of BM significantly fits to the active site of p75NTR. Further Molecular Dynamics simulation study was performed to examine the stability of the binding of these phytochemicals with the selected targets. Our findings suggested that the phytocompounds Bacopasaponin D and Bacopasaponin G significantly binds with p75NTR and thus might have a potential to inhibit the natural binding activity of amyloid-b plaques and act as a potential anti-neurodegenerative drug.

## 1. Introduction

Proper functioning of neurons is crucial for a healthy brain. Alteration in the functioning may produce neurodegenerative disorders (NDs). NDs are recognized to be noteworthy threats to human health. Numerous symptoms are associated with NDs including memory loss, tremors, forgetfulness, agitation. Many kinds of mechanisms lead to NDs such as apoptosis, protein aggregation, oxidative stress, cytotoxicity and ageing. NDs constitute various types of disorders which include Parkinson’s [1], Huntington’s [2], Alzheimer’s disease and other dementias [3], amyotrophic lateral sclerosis [4], frontotemporal dementia [5], spinocerebellar ataxias [6], stroke [7], meningitis [8], encephalitis [9], tetanus [10], epilepsy [11], multiple sclerosis [12], motor neuron disease [13], migraine [14], tension-type headache [15], medication overuse headache [16], brain and nervous system cancers [17], and a residual category of other neurological disorders. There is huge burden of NDs on the global, regional and national levels.

Alzheimer’s disease (AD) has received considerable attention due to its rapid increase in affecting large population. Dementia is one of the common complications occurring as a result of AD, which is responsible for loss of cognition and memory. Excess production of amyloid beta (Aβ) protein is reported to be one of the main contributors to degeneration of neurons in brain. Produced Aβ forms sensile plaques which is deposited in the form of harmful oligomers and initiate AD dementia. Amyloid beta proteins are produced by proteolytic cleavage of amyloid precursor proteins by β-secretase enzyme [18]. It is proved by several studies that Interaction of Aβ42 with a low-affinity growth factor receptor P^75NTR^ actually leads to neuronal loss.

Several synthetic drugs are being used to treat different neurological disorders [19]. for example, Multiple Sclerosis (MS) a neurological disease is treated by wide arrays of repurposed drugs. Some drugs which proved to be effective in mitigating the effect of MS are mitoxantrone [20], cyclophosphamide [21], cladribine [22], amiloride [23], and ibudilast [24]. Use of synthetic drugs is being restricted because of their various kind of side effects such as headache, pain, toxicity, nausea, alopecia, male and women infertility and risk of malignancy. Mitoxantrone is reported to impart many kinds of side effects in the patient suffering from MS [25]. Cyclophosphamide and cladribine also have shown significant side effects such as altered locomotor activity and polyneuropathy [26]. Due to these numerous side effects of synthetic drugs researchers are focused of plant-based drug discovery for the treatment of different neurological disorders. Plants are known to have therapeutic effects pertaining to neurodegenerative disorders [27]. It is revealed that different metabolites present in plants such as phenolics, isoprenoids and alkaloids etc. are mainly responsible for neurotherapeutic actions. Phytochemicals from medicinal plants have been exploited for their potential in targeting neurodegenerative disorders by regulating the Neurotrophins. Example of some plants whose phytochemicals target NDs by affecting Neurotrophins are *Aster scaber* [28], *Camellia sinensis* [29], *Curcuma longa* [30], *Ginkgo biloba* (L) [31], *Liriope platyphylla* [32] and *Magnolia officinalis* [33]. *Bacopa monneiri* (BM) is a significant Ayurvedic medicinal plants that has been in use since ancient time to enhance brain function and to improve intelligence as well as memory [34]. Role of *B. monneiri* in cognitive and memory enhancement has been studied extensively [35]. *B. monneiri* contains many bioactive components such as saponins, triterpenoid bacosaponins and saponins glycosides [35]. It is reported that *B. monneiri* extract enhanced the cognition safely and effectively [36]. Standard extracts of *B. monneiri* have shown improvement in many neurodegenerative disorders owing to presence of different bioactive phytochemicals [33, 36-38]. Aerial parts of *B. monneiri* were found to be cognitive enhancer and neuroprotectant against Alzheimer’s disease [39]. In an in-vitro study Dhanasekaran et al. [40] demonstrated that leaves extract of *B. monneiri* provide neuroprotective mechanism in Alzheimer’s. According to Kumar et al. [41] when extracts of aerial parts of BM administered orally produce neuroprotective effect in cold stress induced hippocampal neurodegeneration of rats.

These fragmented pieces of evidence show the potential of *B. monneiri* to act as a potent anti-ND drug. However, to attain such therapeutic effect using *B. monneiri* phytochemicals is a tedious job as the synthesis of lead phytochemical and their analogs is a time-consuming and economically costly job. Thus, in such a predicament, virtual screening of potential drug candidates is found to be more reliable, focused and cost-effective [42]. Molecular docking and simulation approaches enhance the number of molecules screened and eliminate the urge for in-vitro analysis of the synthetic molecules [43]. With the added advantage that the virtual docking and simulation study provides, this work was carried out to screen the phytochemicals of *B. monneiri* and to assess their potential to be used as a neuroprotective agent against Alzheimer’s. This study could lead to significant advantage in the field of Neurotherapy by development of potential plant-based drug for the treatment of Alzheimer’s disorders.

## 2. Methodology

### 2.1. Phytochemical Selection

*Bacopa monneiri* contains many phytocompounds that belong to the class of saponins, triterpenoids, bacopa saponins and saponins glycosides [35]. Among these phytocompounds, eight phytocompounds were selected.

### 2.2 Ligand and Receptor Preparation

The 3-D structures of selected phytocompounds (sdf format) and their canonical SMILES were downloaded from PubChem Compound (https://pubchem.ncbi.nlm.nih.gov/) database [44] and converted to PDB format using Open Babel software (http://openbabel.org/). The 2D structure of those phytochemicals whose 3D structures were not available in the PubChem database were downloaded and converted into 3D using Open Babel software. The energy minimization was carried out using Chem3D software (ChemOffice 2002) [45] through MM2 minimization.

For preparation of the receptor structures, the PDB structure of p75NTR (PDB ID: 3BUK) receptor was downloaded from RCSB Protein Data Bank (www.rcsb.org). The receptor was prepared by deleting all the water molecules and unwanted residues followed by energy minimization. Chimera [46] was used for the visualization of the 3D structure of receptor and natural ligand (Aβ42) in the present study.

### 2.3. Molecular Docking

The first model of protein data bank entry 1IYT [47] served as the structure of amyloid beta protein. The crystal structure of p75NTR was obtained from protein data bank entry 3BUK [48]. The crystal structure of p75NTR is present in a symmetrical 2:2 ratio with neurotrophin in the mentioned PDB ID. For the current study we performed the docking analysis between a single monomer of p75NTR and Aβ42 oligomer using Cluspro 2.0 [49-52]. The best model was selected based on the cluster size and parameters generated by the server.

Autodock 4.0 [53] software was employed for the molecular docking studies between the receptor (PDB ID: 3BUK) and the ligands (phytochemicals). The active sites of the receptor served as the ligand binding site. The detailed methodology for the protein preparation and docking procedure is given in the Supplementary File. The results were visualized using LigPlot^+^ v.2.2 [54].

### 2.4. Molecular Dynamic Simulations

MD simulations were performed to analyze the binding behavior of the phytochemicals to p75NTR in order to study the dynamic behavior of the protein-ligand interactions. The MD simulation was carried out using CABS-Flex 2.0 server (http://biocomp.chem.uw.edu.pl/CABSflex2) [55] by keeping the default parameters. The root mean square fluctuations (RMSF) were assessed and plotted to equate the flexibility of each residue in the–ligand-protein complexes. The RMSF of the protein ligand complex denoted the minimized fluctuation for all the complexes. The Python package, Matplotlib [56], was used for plotting the RMSF curves.

## 3. Results

### 3.1 Selection of Phytochemicals

8 biologically active phytocompounds were selected for the study that were referred from previous literature (Lal and Lal 2020). The structure of phytochemicals is presented in Fig.1. Their structures, molecular weights were verified from PubChem (https://pubchem.ncbi.nlm.nih.gov/) database [44].

**Fig 1.**
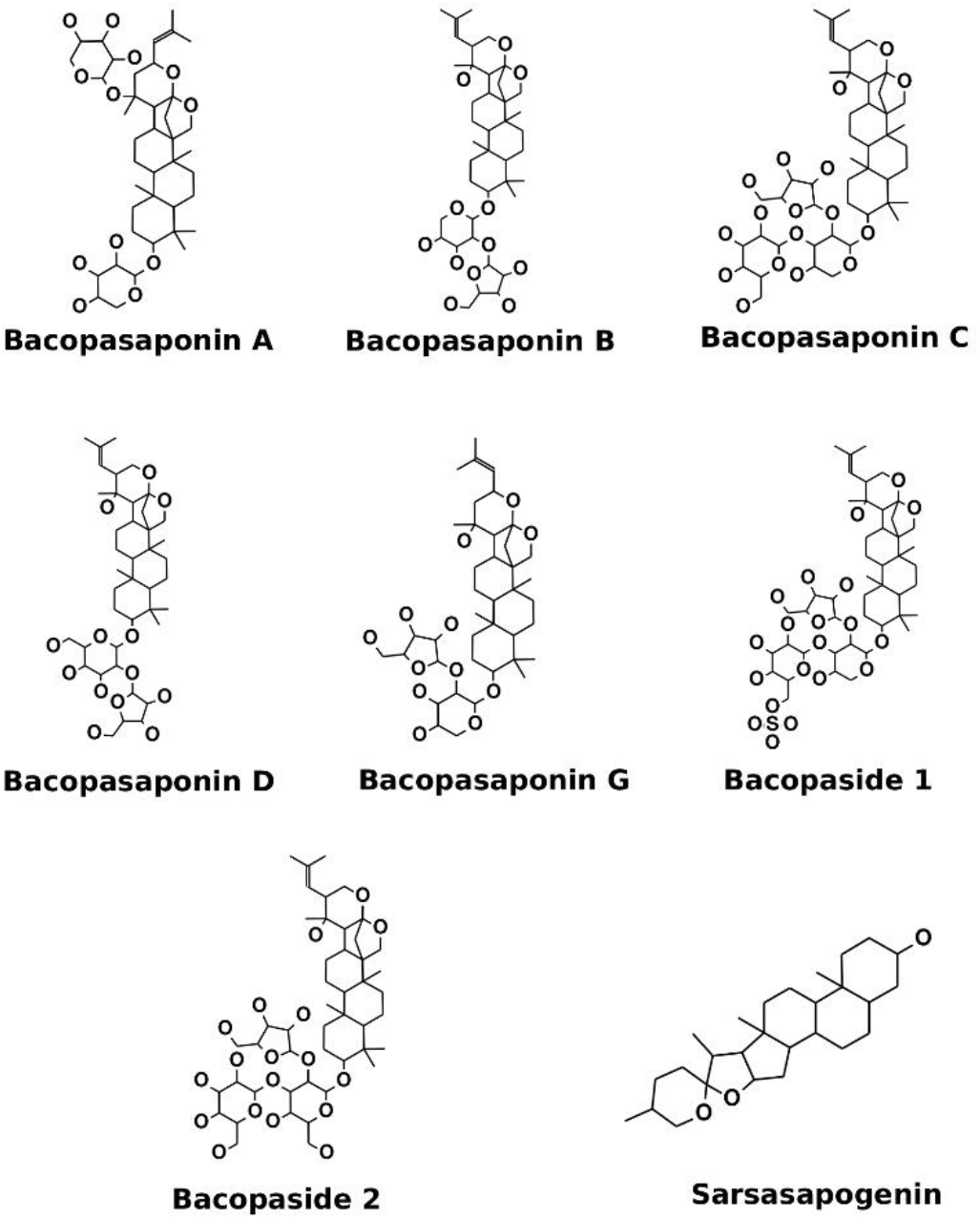
Active Phytochemicals of *Bacopa monneiri*. The structure and molecular weight were verified from PubChem Database

### 3.2. Ligand and Receptor preparation

The interaction between p75NTR and 42 amino acid length Aβ protein was studied to examine the active interaction of amino acid residues. Aβ42 protein monomer and its interaction with p75NTR established the key residues involved in the protein-protein dynamic interaction.

### 3.3. Molecular Docking

Molecular Docking between p75NTR ectodomain monomer and Aβ42 was carried out using Cluspro. Cluster analysis was carried out by the server and 29 clusters were identified by the server with the best cluster having more cluster members and lowest energy. Based on the selection parameters, we selected cluster 2 as it showed lowest energy (−848.7 kcal/mol) and desirable interactions. The anticipated model is acceptable based upon its electrostatic, van der Waals and cluster size. Our docking analysis revealed several hydrogen bond interactions between the ectodomain and the beta amyloid resulting in the stabilization of the complex (Fig.2a). The sequence region between residues 29-35 of Aβ is considered significant for the impact interceded by p75NTR [57]. However, we observed that Ile41 and Gly37 of Aβ42 are hydrogen bonded with Ser5, Thr3 and Thr6 of p75NTR respectively (Fig.2b). We further observed that the Aβ42 favors the topical region of p75NTR commonly termed as the “cap” region [58]. The hydrogen bond interactions between respective proteins are listed in Table 1. These interactions lay down the target site for the small molecules to be developed to target the interaction of p75NTR and beta amyloid proteins.

**Table 1:**
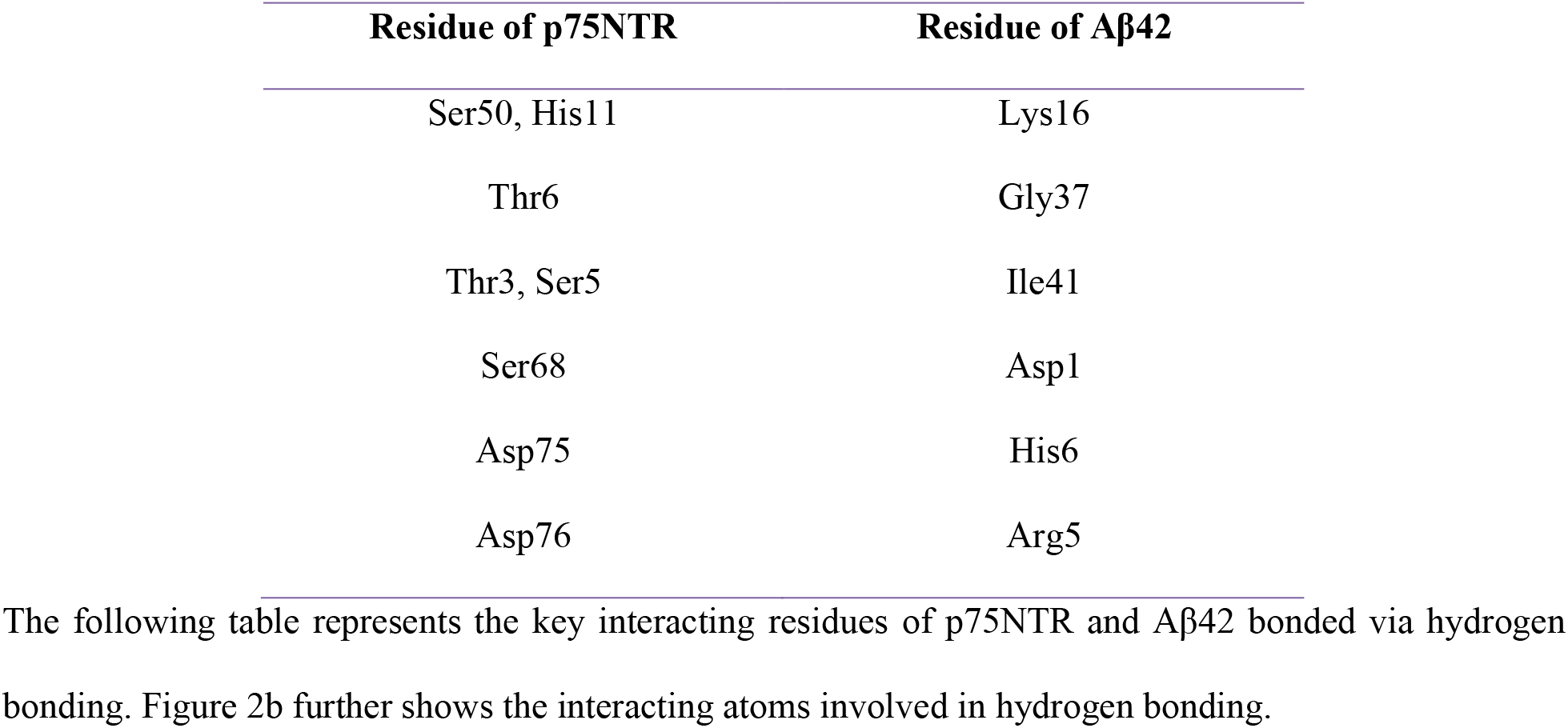
Table representing the hydrogen bonded residues in p75NTR and Aβ42

| Residue of p75NTR | Residue of Aβ42 |
| --- | --- |
| Ser50, His11 | Lys16 |
| Thr6 | Gly37 |
| Thr3, Ser5 | Ile41 |
| Ser68 | Asp1 |
| Asp75 | His6 |
| Asp76 | Arg5 |

**Fig 2.**
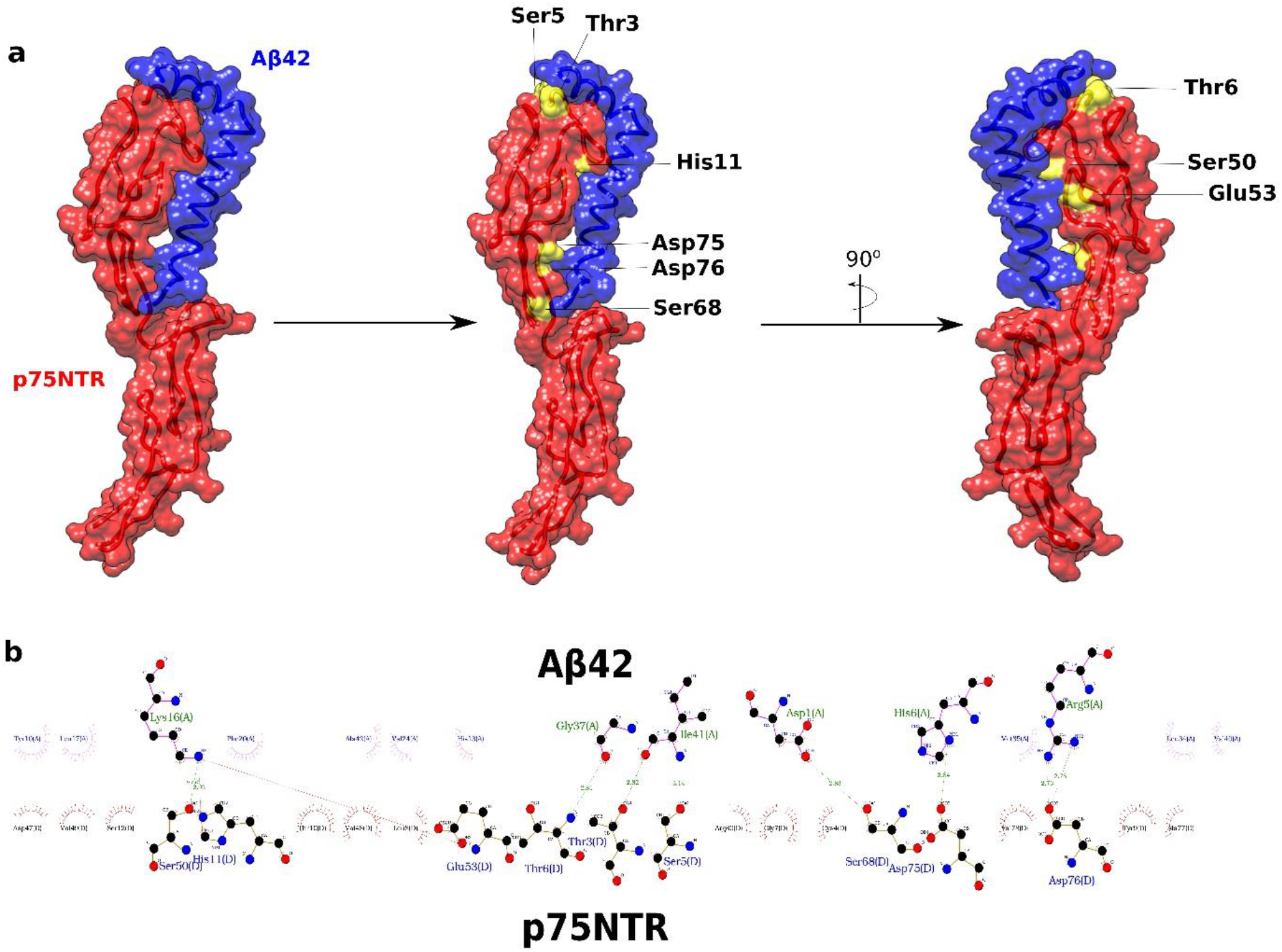
Representation of the active site of the p75NTR and the interaction of p75NTR with amyloid-beta. (a) Molecular Surface representation of the p75NTR receptor (red) in association with amyloid-beta (blue). The key/active residues involved in the binding association are highlighted by yellow color. (b) 2D view of the binding interaction between p75NTR and amyloid-beta protein. The figure shows all the possible interaction occurring between both the molecules with hydrogen bonding being represented by green color

AutoDock 4.0 was employed for the molecular docking studies between the receptor (PDB ID: 3BUK) and the ligands (phytochemicals). The binding affinity between the receptor and ligand was determined based on binding energy value obtained. Out of the eight phytochemicals docked against p75NTR, two phytochemicals, Bacopasaponin D and Bacopasaponin G, showed better interaction with p75NTR. Bacopasaponin D showed hydrophobic interactions with Asp76 along with hydrogen bonding with Cys58 and Val43 with a binding energy of −4.82 kcal/mol (Fig 3a). On the other hand, Bacopasaponin G showed a hydrophobic interaction with Cys79, another important residue of the CRD2 domain of p75NTR involved in the interaction with Aβ42. Apart from hydrophobic interaction, Bacopasaponin G showed hydrogen bond interaction with Asp76, Aps75 and Ala74, which are the key residues involved in the interaction of p75NTR and Aβ42, with a binding energy of −5.39 kcal/mol (Fig 3b). The docking analysis revealed the wide range of interaction of *Bacopa monneiri* phytochemicals with the p75NTR receptor. This analysis is supported by the fact that Aβ42 has a wide range of interaction with p75NTR ranging from recognizing the N-terminal region of the receptor (Fig 2b) to having a binding affinity with the CRD2 domain of p75NTR [58].

**Fig 3.**
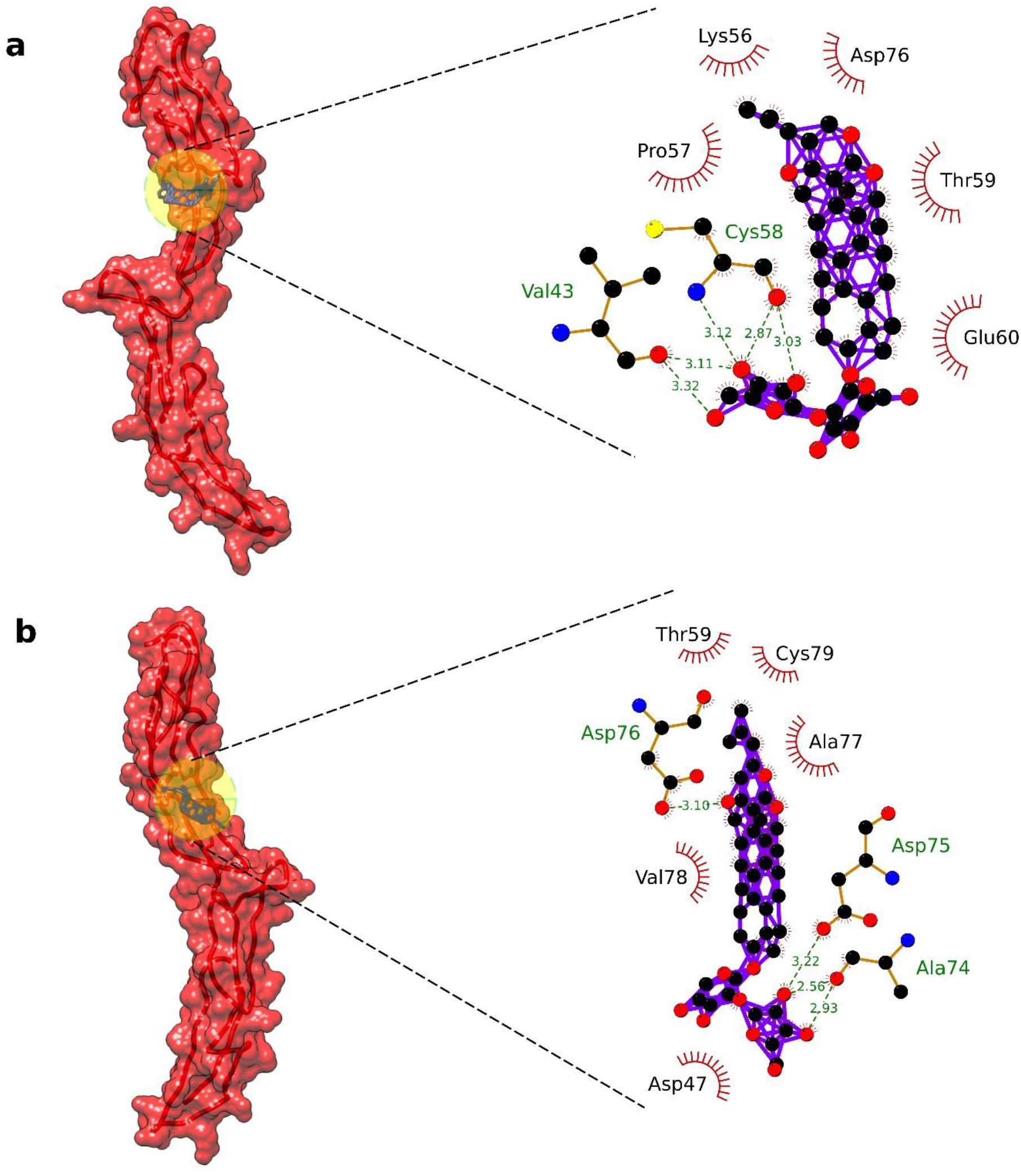
Molecular Docking representation of the top-hit phytochemical of Bacopa monneiri with p75NTR. (a) Docking interaction of Bacopasaponin D with p75NTR showing a hydrogen bonding with Val43 and Cys58. (b) Docking interaction of Bacopasaponin G with p75NTR showing hydrogen bonding with Asp76, Asp78 and Ala74

### 3.4. Molecular Dynamic Simulation

The Molecular Dynamic Simulation was performed using CABS-Flex 2.0 server by keeping the default parameters the same. The RMSF activity of the phytochemicals was observed throughout the simulation as it binds with the p75NTR receptor. The Root Mean Square Fluctuation (RMSF) exhibits the flexibility at residual level. Low level of fluctuations was observed in Bacopasaponin G (Fig.4). The RMSF fluctuation plots reveal the stable behavior of the functionally important residues. The RMSF fluctuation plot of Bacopasaponin G resembles closely to that of Aβ42 (Fig 4). On the other hand, Bacopasaponin D had a high fluctuation in the topical region of p75NTR suggesting that the binding efficiency of the phytochemical is more favored towards the CRD2 domain of p75NTR as compared to the “cap” region. As compared to the other regions of p75NTR, the region ranging from residue 140-150 showed higher fluctuation with both the phytochemicals as well as Aβ42. This fluctuating behavior further suggests the long-range interaction of Aβ42 with the CRD domain of p75NTR. Both the phytochemicals resembled a similar pattern in the fluctuating region suggesting the binding efficiency of the phytochemicals with p75NTR.

**Fig 4.**
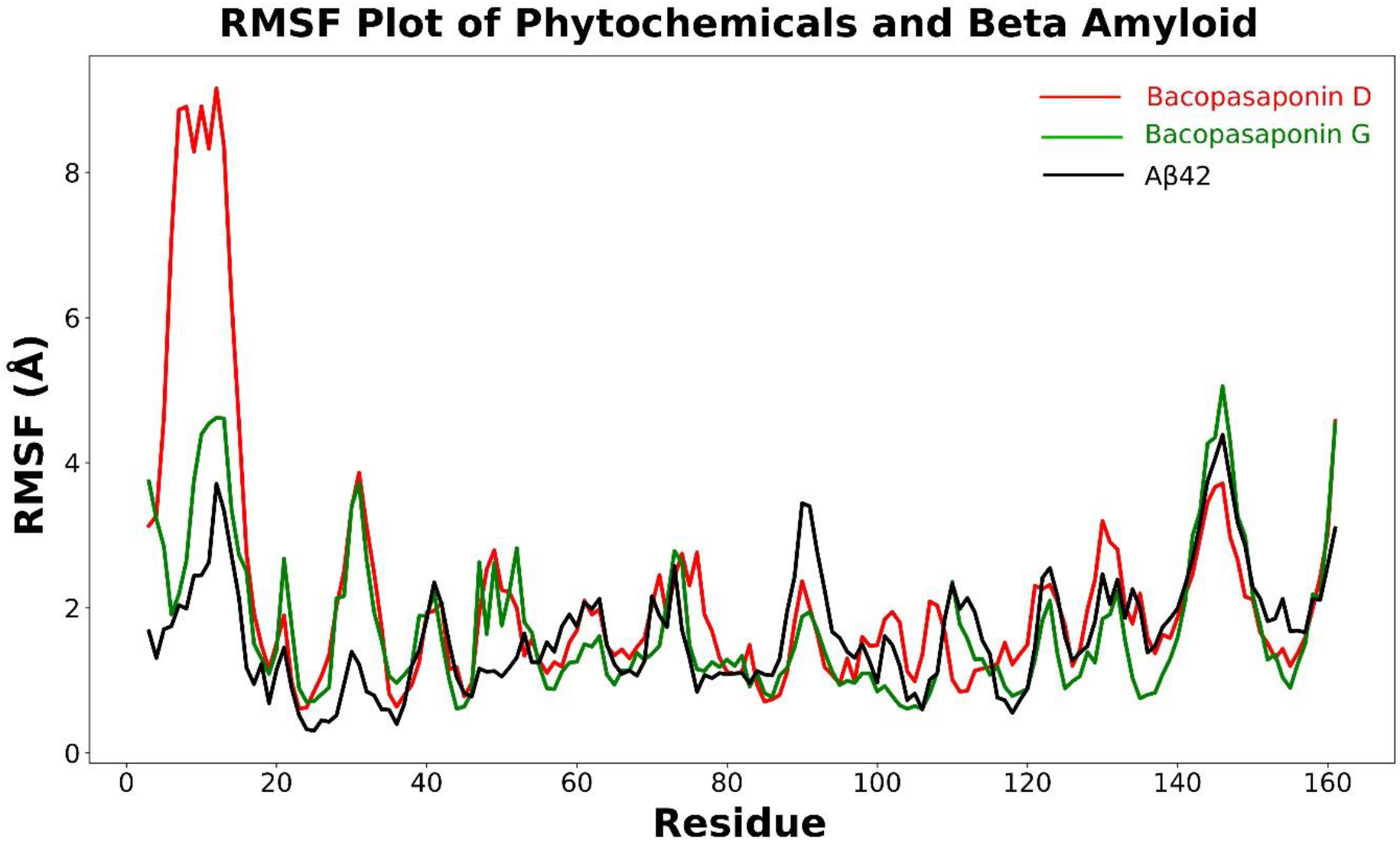
Root Mean Square Fluctuation of the top-hit phytochemical and amyloid-beta with p75NTR. Both Bacopasaponin D and Bacopasaponin G showed a similar behavior as the binding of amyloid-beta with p75NTR. Moreover, Bacopasaponin G had a better RMSF behavior as compared to amyloid-beta whereas Bacopasaponin D favored the CRD2 domain as compared to the topical region of p75NTR

## Conclusion

p75NTR acts as a neurotrophin receptor and thus acts as a potential target for treating neurodegenerative disorders. Aβ42 acts as a natural ligand for p75NTR and its binding results in the nerve cell apoptosis and signal cascade activation [59]. Molecular docking was performed to first investigate the binding characteristics of Aβ42 with p75NTR using Cluspro. The results showed that Aβ42 has a strong binding affinity with p75NTR and this interaction is stabilized by a series of hydrogen bond interactions. Devrajan and Sharmila (2014) have obtained a similar binding behavior of Aβ42 with p75NTR. The investigation related to binding behavior of Aβ42 is also supported experimentally [60-62]. The protein-protein interaction study was followed by the molecular docking and molecular dynamic simulation of phytochemicals of *Bacopa monneiri* with p75NTR with the aim of identifying potential inhibitors. The results showed that two phytochemicals of *Bacopa monneiri*, Bacopasaponin D and Bacopasaponin G, has a good binding affinity towards the active region of p75NTR, suggesting their role as a potential inhibitor in cases of p75NTR induced apoptosis. RMSF Fluctuation shows that Bacopasaponin G indeed has a very close binding characteristic to that of p75NTR and Aβ42 suggesting that it almost mimics the binding behavior. These results provide a good framework to forecast the potential of *Bacopa monneiri* phytochemicals as potent inhibitors. Further investigation in this domain could elucidate the intricate details of these phytochemicals and how they are imparting their activity because p75NTR is being an important therapeutic target, also is a crucial factor between neuronal survival and death in cases of Alzheimer’s.

## Acknowledgement

The authors would like to thank the creators of respective open-source software available to perform the required experiments.

## Competing Interests

The authors declare no completing interests

## Data Availability

The structure of respective phytochemicals used in the study can be freely downloaded from PubChem database [44] and the structure of the proteins can be freely downloaded from Protein Data Bank (PDB) (https://www.rcsb.org/). Cluspro [49-52] is available has an open-source free server for protein-protein docking. AutoDock [53] software is freely available to the user to download from the official website of Scripps Research Institute (https://autodock.scripps.edu/download-autodock4/). CABS-flex 2.0 [55] is freely available as an open-source server (http://biocomp.chem.uw.edu.pl/CABSflex2). LigPlot [54] was used for the visualization.

## Notes

### Competing Interest Statement

The authors have declared no competing interest.

